# Olfactory Bulb Pinprick Induction of Cortical Spreading Depolarizations

**DOI:** 10.1101/2025.09.17.671112

**Authors:** Muhammed Miran Öncel, Michael J. Dora, Lydia Hawley, James H. Lai, Elyssa M. Alber, Joanna Yang, Andrew Thompson, Camila Mera, Roderick Bronson, Andreia Morais, Sava Sakadžić, Andrea Harriott, Cenk Ayata, David Y Chung

## Abstract

Cortical spreading depolarization (CSD) is a wave of cellular depolarization followed by prolonged depression of neuronal activity and is associated with a broad array of neurological diseases, including migraine with aura, traumatic brain injury, and stroke. Traditional CSD induction methods for animal studies have included pinprick, concentrated potassium chloride (KCl) application, and electrical stimulation. These methods are invasive and can cause injury to the cortex. Recently, a non-invasive approach using optogenetics has become available, but requires the use of transgenic mice or transfection of an optogene, which limits its wide adoption. Here, we describe a novel approach using olfactory bulb needle insertion in rodents to induce CSD. We also included KCl-induced CSDs as a comparator in the same mice. Olfactory bulb pinprick resulted in CSDs on every attempt (n = 18/18) as confirmed with optical intrinsic signal imaging. Histological analysis revealed that needle disruption in the caudal olfactory bulb, which is continuous with the cerebral cortex, may account for the propagation of CSD from the olfactory bulb to the cortex. Olfactory bulb pinprick reliably induces CSD and is non-invasive with respect to cortex. The approach may prove to be useful in rodent studies where maintenance of cortical integrity is important.

## Introduction

Cortical spreading depolarization (CSD) is an electrophysiological phenomenon implicated in the pathophysiology of migraine with aura, traumatic brain injury, and stroke^1,2^. Preclinical models are important to determine where CSD lies in causal pathways; however, typical induction methods are invasive and can lead to tissue injury, ischemia, and inflammation which confound the interpretation of experimental results^3-6^.

All existing CSD models involve manipulation of the cortical surface. The 3 historically predominant methods are highly invasive and involve pinprick, application of high concentration potassium chloride, or electrical stimulus through a burr hole^7-11^. Recently, less invasive approaches have been developed. For example, potassium chloride applied to the cortical surface through thinned skull may minimize tissue injury^4,12^. The least invasive approach involves the use of optogenetics which enables cortical induction of CSD with light through intact and unaltered skull^13^. The major limitation of optogenetics, however, is the necessity of either creating a transgenic animal line or an invasive transfection process to introduce the optogene.

Therefore, we sought to develop a novel way to induce CSDs to address the limitations of existing models. We recently observed CSD in the setting of needle insertion through mouse olfactory bulb^14^. Isolated disruption of olfactory bulb can be viewed as non-invasive with respect to cerebral cortex. Here, we refined the olfactory approach and compared it to cortically-based KCI-induced CSD.

## Methods

### Animal Subjects

We adhered to the Animal Research: Reporting of In Vivo Experiments (ARRIVE) guidelines 2.0. All animal protocols were approved by the Institutional Animal Care and Use Committee (Massachusetts General Hospital Subcommittee on Research Animal Care). A total of 21 mice (11 males, 10 females) between the ages of 11 and 13 weeks with an average age of 12.2 ± 0.4 (SD) were used for this study. The mice were housed in groups of 2-4 in a facility accredited by the American Association for Accreditation of Laboratory Animal Care (AAALAC). The animals were kept under diurnal lighting conditions, at a room temperature of 25 °C, and with air humidity ranging from 45% to 65%.

### Experimental Protocol

The mice were placed under anesthesia using isoflurane, with a 5.0% induction dose and a 2.0-2.5% maintenance dose in a gas mixture of 70% N_2_O and 30% O_2_. They were then securely positioned in a stereotaxic frame, and lubricating ointment was applied to protect their eyes. A homeothermic heating pad maintained their body temperature at 36.5°C, monitored via a rectal probe. A midline scalp incision was made to expose the skull overlying the olfactory bulb to the visual cortex. Mineral oil was immediately applied to the skull surface to prevent drying and opacification. The periosteum and other connective tissues were carefully dissected, and mineral oil was continuously applied as necessary. Intraoperative optical imaging (using a USB camera and YawCam software) was initiated at a frequency of 1 Hz and a resolution of 640 × 480 pixels. This was done to identify any inadvertently triggered CSDs during skull preparation, which would serve as an exclusion criterion. A custom MATLAB script was used to confirm the occurrence of CSDs^15^. Subsequently, a burr hole with a 1 mm diameter was drilled over the right olfactory bulb, 5 mm anterior to bregma and 1 mm lateral to the midline, while avoiding the frontal or sagittal venous sinuses to minimize bleeding. A skull thinning procedure^4^ was performed overlying the left cortical surface, 1 mm posterior to the frontal sinus and 2 mm lateral to the midline (Figure 1a). Two animals were excluded from the final analysis as a CSD was accidentally induced during this preparation phase.

**Figure 1.**
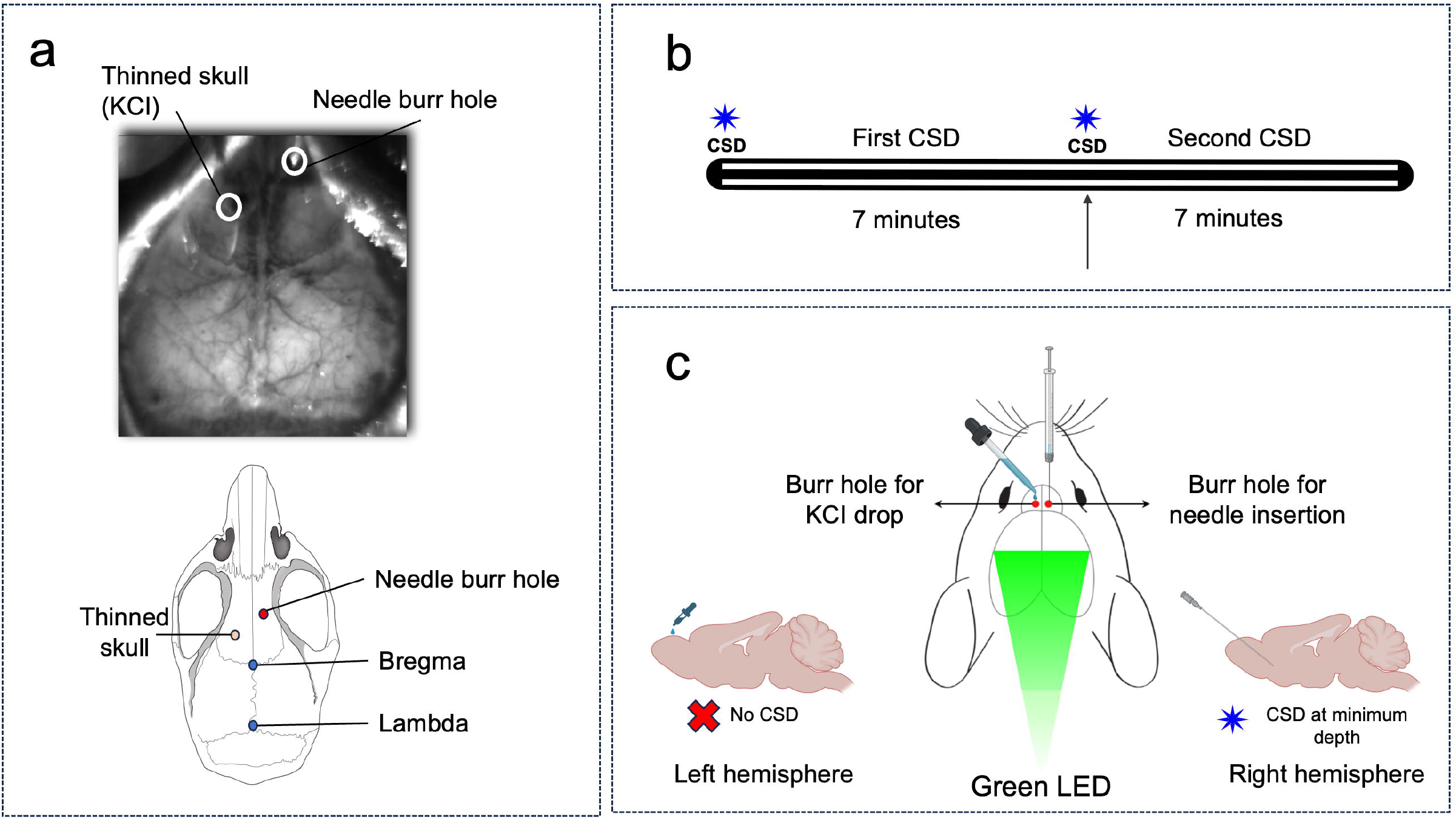
Field of view in intraoperative OIS camera and needle/KCI morphometry **(a).** The timeline shows the experimental protocol **(b)**. Experimental protocol for CSD induction. In half of the animals, the first CSD was induced with KCl. The order of the CSD induction approach was reversed in the other half of the animals. A separate set of experiments was conducted to determine the minimum depth required to induce CSD and to assess whether a CSD can be triggered by applying a KCl drop to the olfactory bulb **(c)**

To control for the potential effect of anesthesia time on CSD characteristics, animals were divided into two age- and sex-matched groups. Briefly, the experiments were conducted as follows: There were 2 CSDs induced for each animal. In half the animals (n=6), a drop of 1M KCl was applied directly on the thinned skull to induce the first CSD in the left hemisphere (An example of the KCl first experiments is shown in Figure 3a.) Once the CSD was visually confirmed, residual KCl was removed with a cotton ball to prevent additional CSDs. After applying KCl, we waited 7 minutes to allow the CSD to complete its transit across the hemisphere (approximately 2 minutes given that a single CSD typically travels at 3–5 mm per minute^7^) and to ensure an adequate refractory period of about 5 minutes. A needle (PrecisionGlide 27G, BD ref# 305109) was then inserted at a 35° angle from the vertical axis (Figure 2b) and parallel to the sagittal plane into the olfactory bulb with the point terminating at the base of the skull. In the other half of the animals (n=6), the procedure was reversed so that needle insertion was used to induce the first CSD in the right olfactory bulb (Figure 1b). A separate set of experiments was conducted to measure the precise minimal depth to cause a CSD and to investigate whether KCI applied on the surface of the olfactory bulb would cause a CSD (n=6) (Figure 1c). Two burr holes, each with a diameter of 1 mm, were drilled over the right and left olfactory bulb, 5 mm anterior to bregma and 1 mm lateral to the midline. A needle was inserted into the right olfactory bulb and advanced 1 mm at a time in 3-minute intervals. The shallowest depth at which the first CSD occurred was noted. Afterwards, 1 M of KCI was dropped onto the left olfactory burr hole, and the mouse was monitored for CSDs. Following the conclusion of the imaging session, the scalp was sutured, and a topical lidocaine ointment was applied.

**Figure 2.**
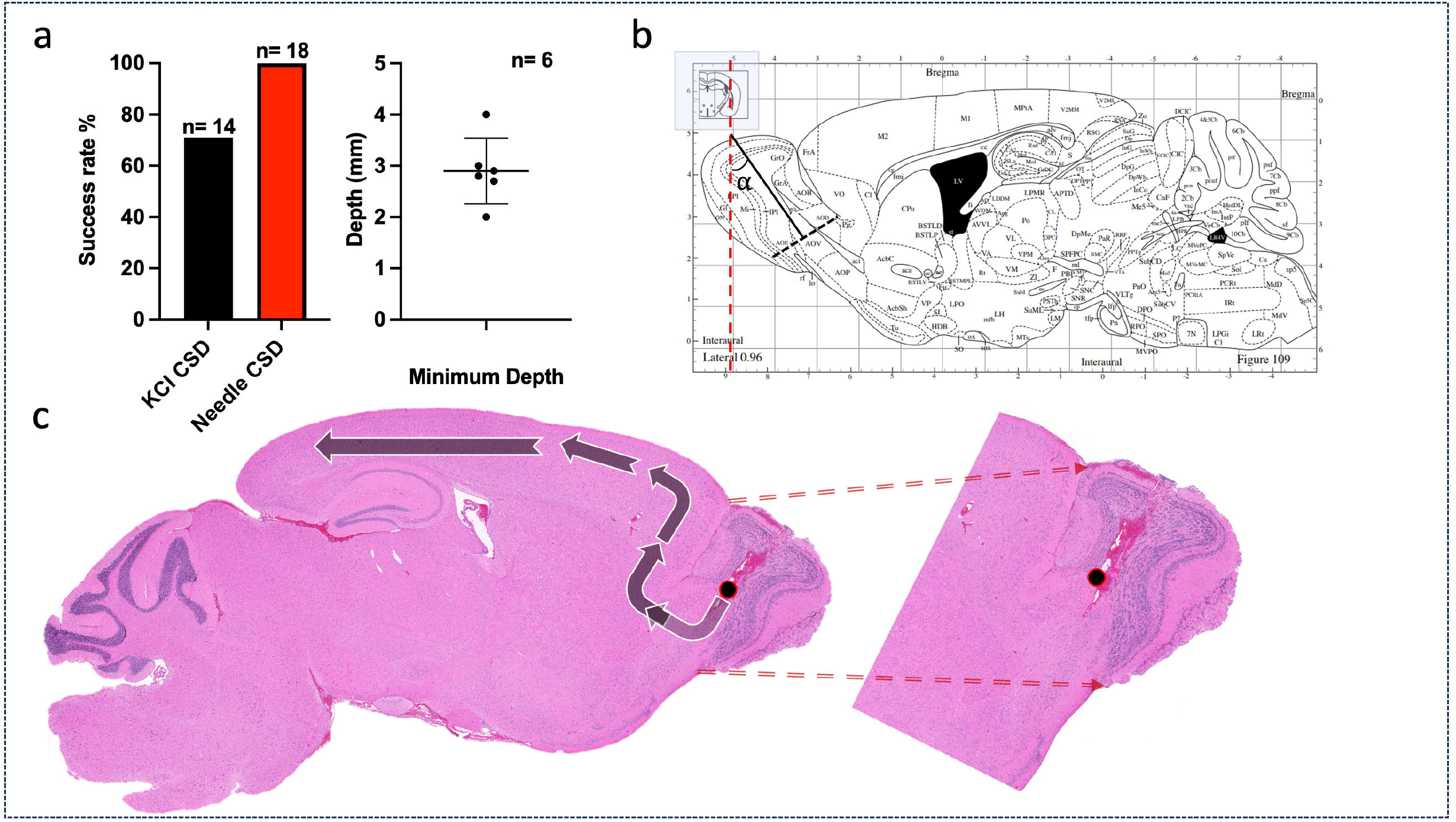
Needle insertion initiates CSD from dorsal olfactory bulb. Minimal depths required to induce a CSD, and success rates of the two approaches **(a).** Midline sagittal mouse atlas with trajectory of needle tract (black line, 3 mm from the cortical surface, (α = 35°)) **(b)**. Sagittal section of an H&E stained brain demonstrates the result of needle insertion and structurally analogous tissue from the tip of the needle tract in dorsal olfactory bulb to the prefrontal cortex **(c)**.

**Figure 3.**
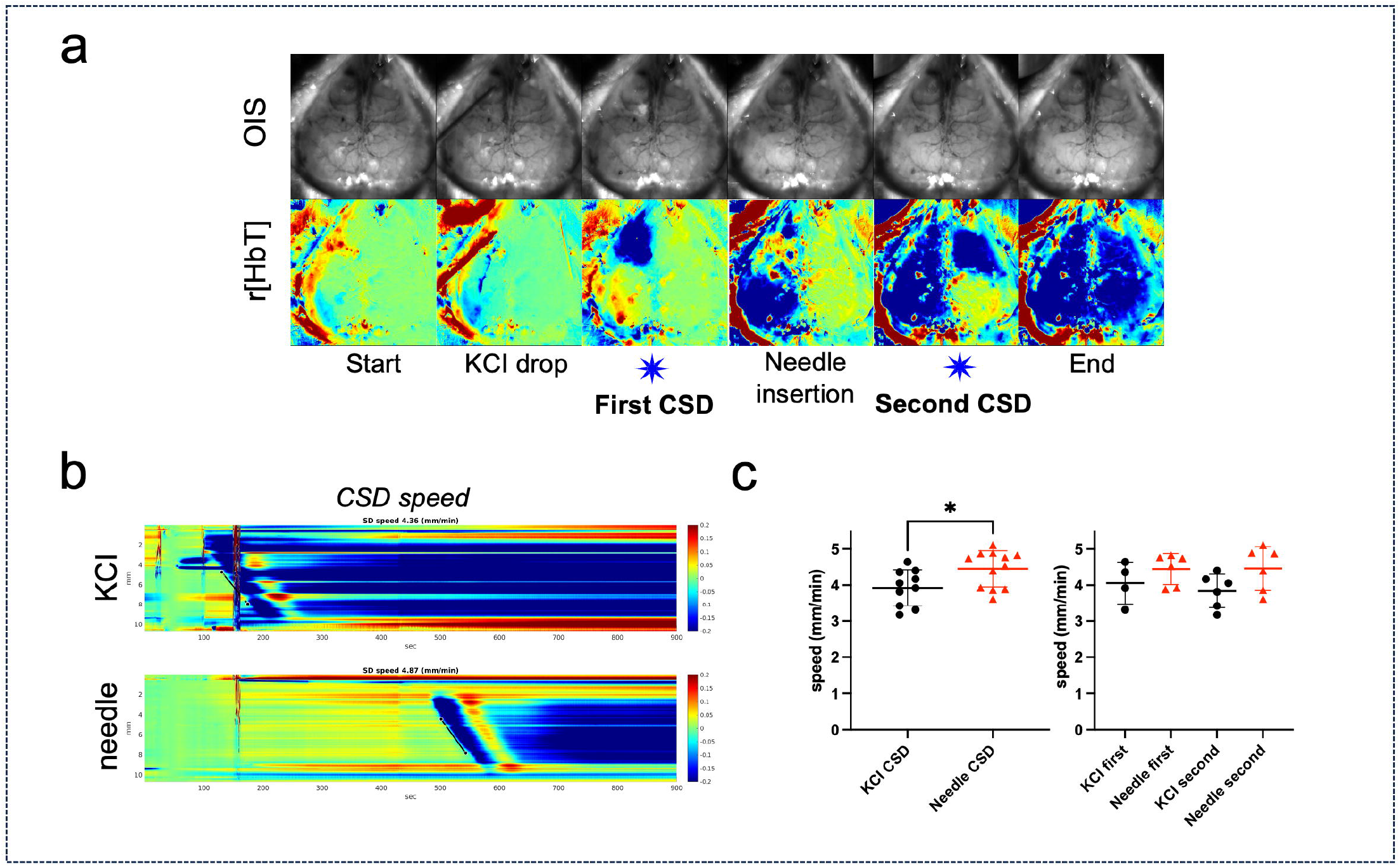
Olfactory needle-induced CSD propagates through cortex faster than KCl-induced CSD. Optical intrinsic signal (OIS) and changes in total hemoglobin r[HbT] from CSDs first induced with KCl (left hemisphere) and then with needle insertion (right hemisphere) **(a).** Total hemoglobin changes in an anterior-posterior line of interest (LOI) **(b)**. The slope of the line across the CSD wavefront is used to calculate the speed of the CSD (mm/min). Group analysis for the speed of the CSD, with time sequence subdivision **(c)**.

### Cerebral Blood Volume (CBV) imaging and data analysis

All of the aforementioned experiments were done while recording cerebral blood volume (CBV) with optical intrinsic signal imaging. The skull surface was diffusely illuminated by a 530 nm green LED (LEDD1B T-Cube LED Driver, M530L3 530 nm Green LED, Thorlabs) and an aspheric condenser lens. Care was taken to minimize surface glare by appropriately positioning the camera and light source. The green channel of the captured images was processed using custom MATLAB code, applying the modified Beer-Lambert law^16^. This processing enhanced the signal changes reflecting total hemoglobin and, thus, CBV. The reflected light intensity of each pixel in each frame was calculated throughout the entire recording period. Two lines of interest (LOIs) were placed, one for the right hemisphere and another symmetrically on the left hemisphere. The speed of the CSDs were calculated by determining the stabilized slope of the wavefront on the LOI r[HbT] (relative total hemoglobin) time course plots (Figure 3b). Prism 10 (GraphPad Software, San Diego, CA, USA) was used for the paired t-test and one-way ANOVA to compare the speed of the CSD for KCI vs. needle and subgroups, respectively. The threshold for statistical significance was a p-value of < 0.05.

### Histology

After a 24-hour post-recording period to allow for inflammatory cell migration, the mice were euthanized under 5% isoflurane anesthesia, and their brains were harvested. The brains were preserved in 4% paraformaldehyde (PFA) for 48 hours and transferred to 1X phosphate-buffered saline (PBS) until sagittal sectioning and staining. The PBS solution was changed weekly. Right-sided sections were easily identified by the needle insertion point. In contrast, visualizing left-sided sections was more challenging. To overcome this, we prepared sagittal sections approximately 1 mm lateral to the midline on the left side to estimate the location of the thinned skull and KCl application site. Hematoxylin and eosin (H&E) stain was used to visualize the needle tract (Figure 2c).

## Results

The duration of the experiments (averaging 51.5 ± 6.7 minutes) was minimized to eliminate the potential confounding effects of anesthesia. CSDs were observed for every needle insertion (n = 18), but in two animals, KCI application failed to induce CSD. Success rates for KCI vs needle induction of CSD were 71% and 100% respectively (Figure 2a). Additionally, intraoperative OIS imaging revealed CSDs during the skull thinning process for two animals; they were excluded from the analysis. KCI drop and subsequent cleaning of the cortical surface introduced noise signals to the r[HbT] data, as visualized in Figures 3a and 3b. The average minimum depth required to induce a CSD using the needle insertion method was 2.9 mm ± 0.6 (n=6) from the brain surface (Figure 2a). We also attempted to induce olfactory CSDs (n=6) by applying KCI to the surface of the olfactory bulb through a burrhole, but no CSDs were observed (Figure 1c). An experienced rodent histopathologist reviewed the H&E slides. Out of 12 provided brains, 4 contained sufficiently preserved sections showing a clear needle tract (Figure 2c), while the remaining slides had sections that missed the course of the needle tract or had tissue preservation or sectioning artifacts. In the evaluable slides, there was evidence of blood extravasation on the needle insertion side, which intersected with the caudal portion of the olfactory bulb in tissue that was histologically contiguous with the cortex on H&E staining (Figure 2c). In contrast, there was no significant tissue disruption apparent on the KCl side in any of the 12 slides.

For the calculation of CSD propagation speed experiments, the needle was inserted at an average speed of 1.0 ± 0.19 mm/sec in order to reliably induce a CSD. The average needle insertion distance from the surface of the brain was 6.27 ± 0.33 mm. A high needle insertion speed is needed to induce CSD by pinprick, since slower insertions may not sufficiently depolarize tissue to trigger the wave^7^. KCl-induced CSD propagation was significantly slower than needle-induced CSD propagation (3.9 ± 0.5 vs. 4.5 ± 0.5 mm/min, p = 0.02; Figure 3c). To assess whether the sequence of CSD induction affected propagation speed, we compared the CSD propagation speeds between the animals that received KCl first versus those that underwent needle insertion first. No significant differences were observed in propagation speed between the two groups (p = 0.1483; Figure 3c).

## Discussion

Preclinical models of CSDs are essential for testing therapies directed at acute brain injury and migraine with aura, yet most available induction methods are invasive^7^. Optogenetic strategies offer a non-invasive alternative but depend on transgenic mouse lines or AAV injection into cortical tissue. Here, we find that olfactory bulb needle insertion induces CSDs with greater reliability than application of high concentration KCl to the cortical surface. Moreover, we identify the dorsal olfactory bulb as the key anatomical pathway through which CSD propagates from the needle insertion site to cerebral cortex.

The pharmacology and molecular mechanisms underlying different CSD induction methods have been reviewed extensively^7^. However, reports specifically describing olfactory bulb induction are limited. To our knowledge, the first description came from a rat study in which potassium injection in the olfactory bulb only occasionally caused CSDs.^17^ Notably, the injections in that study were rescricted to a depth of 1 mm. By contrast, in our prior work in mice, deep needle insertion through the olfactory bulb as part of an injury model invariably triggered CSDs.^14^ The present study builds on that observation, showing that an average depth of 2.9 mm is required to reliably induce a CSD, thereby localizing the site of origin more precisely.

Our findings also revealed that KCl-induced CSD exhibited a slower propagation speed compared to needle-induced CSD (Figure 3c). This difference may be due to molecular changes occurring in the microenvironment specific to each CSD modality. For example, voltage-gated Ca^2+^ (Cav) channels contribute substantially to CSD triggered by high-dose KCl application but are less involved in pinprick-induced CSD^18^. Conversely, the inhibition of the voltage-gated Na^+^ (Nav) channels blocks CSD caused by pinprick^19^.

In conclusion, induction of CSD through the mouse olfactory bulb is a reliable and minimally invasive method that eliminates the need to manipulate the cerebral cortex. The dorsal olfactory bulb acts as a bridge that enables CSD propagation from the olfactory bulb to the prefrontal cortex. This method is advantageous for CSD induction due to its innate nature of minimal cortex disruption, especially for experiments that require cortical integrity.

## Data access

The code used in this paper has already been published and is available to the public^15^. Further data is available upon request to the corresponding author (DYC).

## Acknowledgments

We thank Ms. Dilara Bahadir for her valuable contributions to this work.

---

**Video 1**. Time-lapse shows operative view (right), and corresponding changes in total hemoglobin (r[HbT]) (left).

## References

1. Ayata C, Lauritzen M. Spreading Depression, Spreading Depolarizations, and the Cerebral Vasculature. Physiological reviews 2015;95(3):953–93. DOI: 10.1152/physrev.00027.2014.

2. Leão A. Spreading depression of activity in the cerebral cortex. Journal of Neurophysiology 1944;7(6):359–390.

3. Ayata C, Shin HK, Salomone S, et al. Pronounced hypoperfusion during spreading depression in mouse cortex. Journal of cerebral blood flow and metabolism : official journal of the International Society of Cerebral Blood Flow and Metabolism 2004;24(10):1172–82. DOI: 10.1097/01.WCB.0000137057.92786.F3.

4. Takizawa T, Qin T, Lopes de Morais A, et al. Non-invasively triggered spreading depolarizations induce a rapid pro-inflammatory response in cerebral cortex. Journal of cerebral blood flow and metabolism : official journal of the International Society of Cerebral Blood Flow and Metabolism 2020;40(5):1117–1131. DOI: 10.1177/0271678X19859381.

5. Bouley J, Chung DY, Ayata C, Brown RH, Jr., Henninger N. Cortical Spreading Depression Denotes Concussion Injury. J Neurotrauma 2018. DOI: 10.1089/neu.2018.5844.

6. Lauritzen M, Dreier JP, Fabricius M, Hartings JA, Graf R, Strong AJ. Clinical relevance of cortical spreading depression in neurological disorders: migraine, malignant stroke, subarachnoid and intracranial hemorrhage, and traumatic brain injury. Journal of cerebral blood flow and metabolism : official journal of the International Society of Cerebral Blood Flow and Metabolism 2011;31(1):17–35. DOI: 10.1038/jcbfm.2010.191.

7. Pietrobon D, Moskowitz MA. Chaos and commotion in the wake of cortical spreading depression and spreading depolarizations. Nature reviews Neuroscience 2014;15(6):379–93. DOI: 10.1038/nrn3770.

8. Zhao HT, Tuohy MC, Chow D, et al. Neurovascular dynamics of repeated cortical spreading depolarizations after acute brain injury. Cell Rep 2021;37(1):109794. DOI: 10.1016/j.celrep.2021.109794.

9. Longden TA, Dabertrand F, Koide M, et al. Capillary K(+)-sensing initiates retrograde hyperpolarization to increase local cerebral blood flow. Nature neuroscience 2017;20(5):717–726. DOI: 10.1038/nn.4533.

10. Chang JC, Shook LL, Biag J, et al. Biphasic direct current shift, haemoglobin desaturation and neurovascular uncoupling in cortical spreading depression. Brain : a journal of neurology 2010;133(Pt 4):996–1012. DOI: 10.1093/brain/awp338.

11. Ayata C. Pearls and pitfalls in experimental models of spreading depression. Cephalalgia : an international journal of headache 2013;33(8):604–13. DOI: 10.1177/0333102412470216.

12. Tamim I, Chung DY, de Morais AL, et al. Spreading depression as an innate antiseizure mechanism. Nat Commun 2021;12(1):2206. DOI: 10.1038/s41467-021-22464-x.

13. Chung DY, Sadeghian H, Qin T, et al. Determinants of Optogenetic Cortical Spreading Depolarizations. Cereb Cortex 2018. DOI: 10.1093/cercor/bhy021.

14. Lai JH, Qin T, Sakadzic S, Ayata C, Chung DY. Cortical Spreading Depolarizations in a Mouse Model of Subarachnoid Hemorrhage. Neurocrit Care 2022;37(Suppl 1):123–132. DOI: 10.1007/s12028-021-01397-9.

15. Chung DY, Sugimoto K, Fischer P, et al. Real-time non-invasive in vivo visible light detection of cortical spreading depolarizations in mice. Journal of neuroscience methods 2018. DOI: 10.1016/j.jneumeth.2018.09.001.

16. Ma Y, Shaik MA, Kim SH, et al. Wide-field optical mapping of neural activity and brain haemodynamics: considerations and novel approaches. Philos Trans R Soc Lond B Biol Sci 2016;371(1705). DOI: 10.1098/rstb.2015.0360.

17. Amemori T, Gorelova NA, Bures J. Spreading depression in the olfactory bulb of rats: reliable initiation and boundaries of propagation. Neuroscience 1987;22(1):29–36. DOI: 10.1016/0306-4522(87)90195-3.

18. Richter F, Ebersberger A, Schaible H-G. Blockade of voltage-gated calcium channels in rat inhibits repetitive cortical spreading depression. Neuroscience letters 2002;334(2):123–126.

19. Akerman S, Holland PR, Goadsby PJ. Mechanically-induced cortical spreading depression associated regional cerebral blood flow changes are blocked by Na+ ion channel blockade. Brain research 2008;1229:27–36.

